# Phagocytes as Plaque Catalysts: Human Macrophages Actively Generate Pathogenic Aβ42 Fibrils with Seeding and Cross-Seeding Potency

**DOI:** 10.1101/2025.10.11.681781

**Authors:** Katerina Konstantoulea, Meine Ramakers, Sarah C. Borrie, Dries T’syen, Daan Moechars, Malgorzata A. Sliwinska, Brajabandhu Pradhan, Giulia Albertini, Grigoria Tsaka, Bert Houben, Mark Fiers, Dewilde Maarten, Dietmar Rudolf Thal, Willem Michael, Jonas J. Neher, Bart De Strooper, Frederic Rousseau, Joost Schymkowitz

## Abstract

The prevailing view frames microglia and macrophages as guardians against amyloid beta (Aβ) accumulation in Alzheimer’s disease (AD). Here, we overturn this paradigm by demonstrating that human phagocytic cells—including differentiated THP-1 macrophages and iPSC-derived microglia—are not merely passive responders but active producers of extracellular, seeding-competent Aβ42 fibrils, the amyloid species most strongly linked to parenchymal plaque formation and neurodegeneration. These cell-generated aggregates differ structurally and functionally from synthetic fibrils, exhibiting heightened seeding activity and the ability to cross-seed tau aggregation, a key driver of AD progression. Notably, Aβ42 fibril formation in this system requires active cellular processes and is exacerbated by loss of TREM2, a major AD risk gene. Transcriptomic profiling reveals an early inflammatory response resembling microglial states observed in human AD models, positioning this system as a tractable, human-relevant platform to dissect the interplay between Aβ aggregation, innate immunity, and genetic susceptibility. Our findings suggest that macrophages and microglia play a dual role in AD, acting both as responders and inadvertent catalysts of pathogenic amyloid formation, with implications for early therapeutic intervention.

**Significance Statement:** How amyloid plaques emerge and spread in Alzheimer’s disease remains a critical unanswered question. Here, we show that human immune cells—including brain-resident microglia—can actively generate Aβ42 fibrils, the form of amyloid most strongly linked to neurodegeneration. These cell-produced fibrils not only seed further amyloid buildup but also trigger tau aggregation, a key event in disease progression. We further demonstrate that genetic risk factors like TREM2 amplify this process. Our findings reveal a direct link between immune cell activity, genetic susceptibility, and the earliest stages of Alzheimer’s pathology, offering new insights into disease mechanisms and potential intervention points.

## Background

Alzheimer’s disease (AD) is a progressive neurodegenerative disorder that remains a major global health challenge due to its increasing prevalence ^1,2^. Central to AD pathology is the accumulation of amyloid beta (Aβ) plaques, particularly fibrils composed of Aβ42, a highly aggregation-prone peptide^3^. The enhanced aggregation and seeding potential of Aβ42 compared to Aβ40 underlie its strong association with disease progression^4,5^, including its role in initiating tau pathology and driving neuroinflammation^6^. Current amyloid therapies targeting Aβ aggregates represent a significant advancement in the treatment of AD. To date, three monoclonal antibodies have received FDA approval for this purpose. Aducanumab, derived from memory B cells of cognitively normal elderly individuals, binds parenchymal Aβ and has been shown to reduce both amyloid plaque burden and cognitive decline^7,8^. Lecanemab, developed from the mouse antibody mAb158, targets soluble Aβ aggregates and has demonstrated efficacy in slowing clinical progression and clearing Aβ deposits^7,9,10^. Donanemab, based on the mouse antibody mE8-IgG2a, targets explicitly the pyroglutamate-modified N-terminus of Aβ—a form enriched in amyloid plaques—and has shown benefits in individuals with early symptomatic AD or mild cognitive impairment^11–13^. Despite extensive research in Aβ pathology and therapeutic development, critical gaps remain in understanding the cellular mechanisms that regulate Aβ aggregation and their contribution to AD progression.

Microglia, the brain’s resident immune cells, are increasingly recognized as central players in the formation and propagation of Aβ plaques. They exhibit a dual role: on the one hand, microglia clear Aβ aggregates through phagocytosis^14,15^; on the other they can promote aggregation and contribute to amyloid seeding^16–18^. For example, depletion of the microglial niche in a mouse of Aβ pathology led to a reduction in plaque load^19^. The opposing functions of microglia are implicated in the early stages of AD, where immune dysregulation may shift microglial activity toward amyloid propagation and inflammation^20^. Understanding how microglia interact with Aβ42 and their role in plaque formation is crucial for unraveling the complex interplay between amyloid deposition and neuroinflammation.

In vitro models that capture microglial behavior are essential tools for mechanistic studies of amyloid aggregation and innate immune responses. THP-1 macrophages, a human monocytic cell line, have previously been shown to support amyloid formation, particularly with Aβ40, a peptide more commonly associated with cerebral amyloid angiopathy and vascular pathology^21,22^. In contrast, Aβ42 is more directly implicated in parenchymal plaque formation and neurodegeneration^4,5^, making it a more relevant target for modeling AD pathophysiology. Studying Aβ42 aggregation in vitro therefore presents a more disease-relevant but also more challenging scenario. Moreover, although macrophages and microglia share phagocytic functions, they exist in distinct anatomical and immunological niches^23^. As such, it cannot be assumed that macrophage cell lines will faithfully replicate microglial behavior, particularly in the context of Aβ42 aggregation and associated inflammatory responses.

TREM2 (Triggering Receptor Expressed on Myeloid Cells 2) has emerged as a critical regulator of microglial function in AD. Genetic variants in TREM2, such as the R47H substitution, are among the strongest risk factors for late-onset sporadic AD after APOE^24–26^. These variants are associated with increased amyloid burden and tau pathology, and loss of TREM2 function impairs microglial ability to respond to plaques^25,27^. Functionally, TREM2 mediates microglial lipid sensing, cell survival, and transition to disease-associated states that promote plaque compaction and modulate neurodegeneration^28^. These insights emphasize the need for in vitro models capable of capturing TREM2-regulated microglial dynamics, a gap this study seeks to fill.

In light of these challenges, a key goal is to establish in vitro models that not only replicate microglial responses to Aβ42 but also respond to AD-relevant genetic perturbations. Here, we show that THP-1 macrophages not only promote the extracellular deposition of Aβ42 fibrils with distinct structural and functional properties, but also recapitulate key features of early microglial responses, including cytokine-driven inflammation^29^ and sensitivity to TREM2 perturbation^30^. Using transcriptomic comparisons with hESC-derived microglia and genetic perturbations in both THP-1 and EPSC-derived microglia, we demonstrate that THP-1 cells represent a tractable and biologically relevant system for studying amyloid-associated innate immune processes. Our findings provide a framework for understanding the role of macrophages in amyloid aggregation and inflammation and establish a relevant model for studying early-stage mechanisms in AD and for evaluating therapeutic interventions.

## Results

### THP-1 macrophages induce extracellular amyloid deposition of Aβ42

To investigate the potential for THP-1 macrophages to deposit Aβ42 aggregates, we differentiated THP-1 cells into macrophages using 50 ng/mL PMA for 48 hours. The differentiated macrophages were then incubated with freshly dissolved Aβ42 at various concentrations for an additional 48 hours (Figure 1A). Initially, we tested deposition in the presence and absence of 100 μg/mL (∼22 μM) Aβ42. Staining with Congo red, the gold-standard dye used for diagnosing amyloidosis due to its apple-green birefringence under polarized light^31^, revealed the expected birefringence in the presence of 100 μg/mL Aβ42, indicating amyloid formation and deposition (Figure 1B). No birefringence was observed in control samples lacking Aβ42 (Figure 1B).

**Figure 1:**
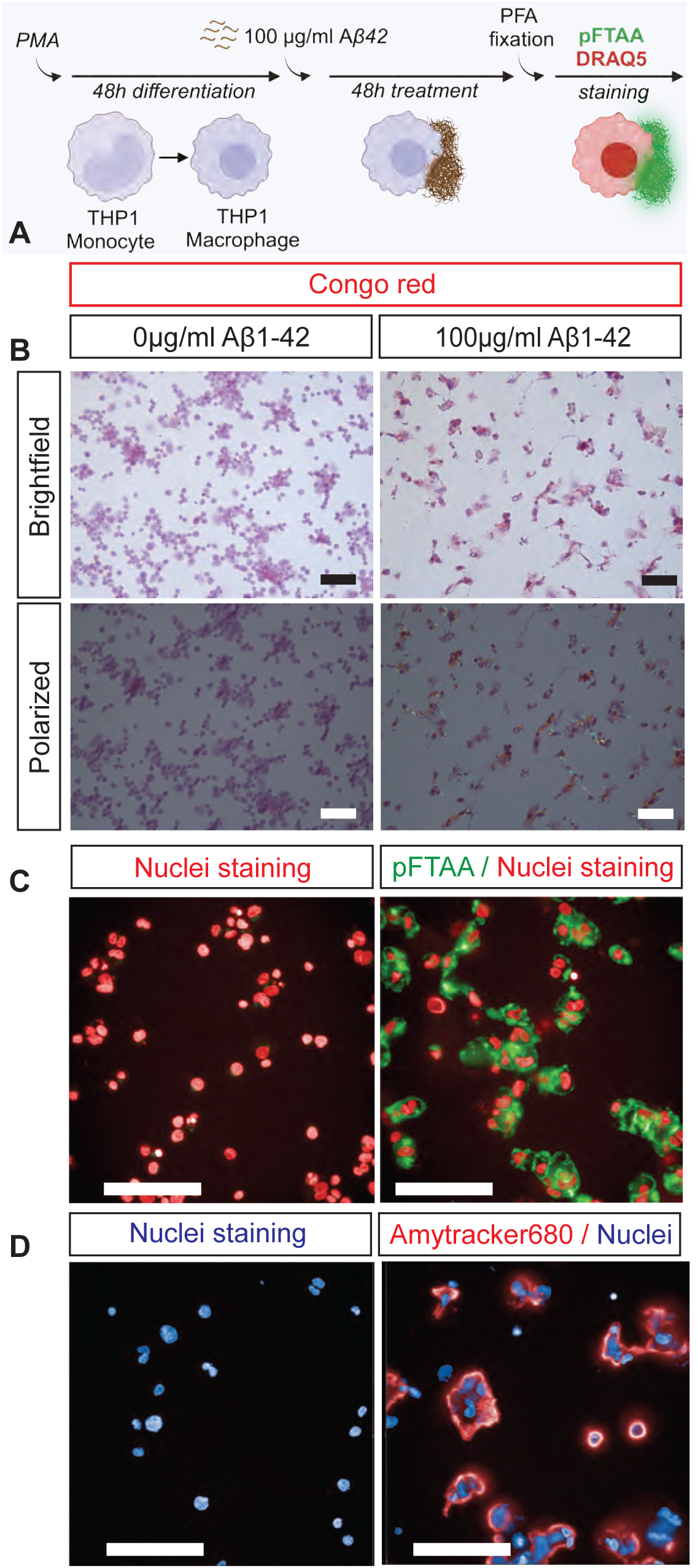
THP-1 cells deposit amyloid fibrils. A) Schematic of experimental procedure B) Congo Red staining in the absence or presence of rAβ42 (0, 100 μg/ml). The panels above are the cells under Brightfield. The panels below are the cells under polarized light. Light green-blue and yellow color indicates amyloid formation. Purple color: hematoxylin. Red: Congo red. Scale bar: 50μM C) pFTAA staining in the presence and absence of rAβ42 (0, 100μg/ml). Green: pFTAA, red: nuclei staining. Scale bar: 100μm D) Amytracker680 staining in the presence and absence of rAβ42 (0, 100μg/ml). Red: amytracker, blue: nuclei staining. Scale bar: 100μm

To further confirm amyloid deposition, we stained the samples with pFTAA^32^ and Amytracker680, two fluorescent dyes specific to amyloid-rich structures. Both dyes exhibited strong fluorescence signals in the presence of Aβ42, confirming the presence of amyloid aggregates (Figure 1C, D). Importantly, no deposition was observed in the absence of cells, indicating that they are necessary for Aβ42 deposition (Supplementary Figure 1A).

Next, we examined Aβ42 deposition across a range of concentrations (0–100 μg/mL). Differentiated macrophages were treated with increasing concentrations of Aβ42, followed by pFTAA staining. Quantification of fluorescence intensity revealed a dose-dependent increase in Aβ42 deposition, with significant accumulation detected at concentrations above 30 μg/mL (Figure 2A, B). Quantifying the surface area stained by pFTAA gave similar results (Supplementary Figure 1C).

**Figure 2:**
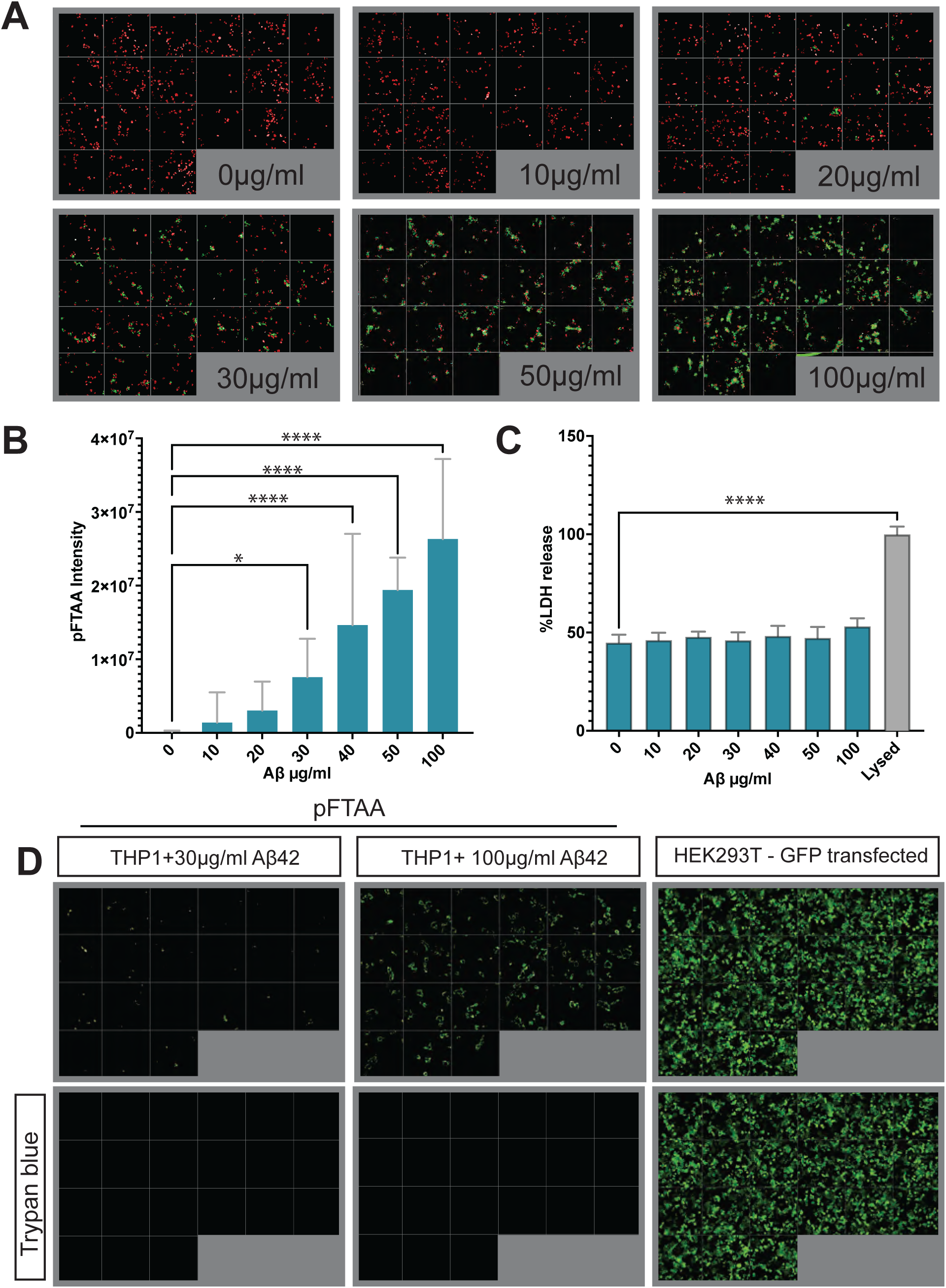
THP-1 cells have significant rAβ42 extracellular deposition. A) Images of Aβ deposition in 7 different rAβ42 concentrations. 21 different fields are shown for each concentration. Red: nuclei (DAPI), Green: aggregates (pFTAA). B) Bar plot of pFTAA intensity of THP-1 cells with increasing concentration of rAβ. 4 independent experiments with 4 repeats, mean+SD. Statistics: Ordinary one-way ANOVA with Dunnett correction for multiple comparisons. *<0.05, ****<0.0001 C) %LDH release in the presence of increasing concentrations of rAβ42. 100% indicates release of lysed cells. 2 independent experiments with 2 repeats, mean+SD. Statistics: Ordinary one-way ANOVA with Dunnett correction for multiple comparisons. ****<0.0001 D) Live imaging of THP-1 cells incubated with 30μg/ml or 100μg/ml rAβ42 and stained with pFTAA (upper panels, left and middle). Lower panels show same wells after addition of Trypan Blue (lower panel, left and middle). Right upper panel shows live imaging of HEK293T cells transfected with GFP. Right lower panel shows same well after addition of Trypan Blue, indicating that Trypan blue successfully quenches extracellular pFTAA staining.

To determine whether the aggregates formed in this model were primarily accumulating in the extracellular space, we employed the Trypan blue quenching technique. This method utilizes Trypan blue’s ability to absorb green fluorescence and its exclusion from live cells. Live cells stained with pFTAA were imaged before and after Trypan blue addition. At both 30 μg/mL and 100 μg/mL Aβ42, most fluorescence was quenched, confirming that the majority of aggregates were extracellular at the time of detection (Figure 2D, left and middle panels). As a control, GFP-expressing HEK cells were similarly treated, and GFP fluorescence persisted after Trypan blue addition, confirming intracellular localization of GFP (Figure 2D, right panels).

Finally, we assessed cell viability using an LDH release assay, which detects membrane disruption. No significant reduction in cell viability was observed, even at 100 μg/mL Aβ42, though a slight but non-significant increase in LDH release was noted at this concentration (Figure 2C). This finding was supported by nuclear counts, which showed no differences in cell numbers after Aβ42 treatment (Supplementary Figure 1B). Interestingly, treated macrophages displayed morphological changes, likely reflecting their activation and capacity to respond to Aβ42, consistent with macrophage plasticity in different activation states.

### Aggregation-prone Aβ42 and active macrophages are essential for amyloid deposition

To verify that the observed deposition of THP-1 cells results from Aβ42 aggregation, we incubated differentiated THP-1 cells with a non-aggregating Aβ42 variant (Scrambled-Aβ42). Scrambled-Aβ42 contains the same amino acids as Aβ42 but has no aggregation propensity as predicted by the TANGO algorithm ^33^ (Supplementary Figure 2A). Additionally, Thioflavin T (ThT), a dye frequently used to monitor aggregation^34^, did not detect amyloid formation of Scrambled-Aβ42 in the concentrations used (10,20,30,40,50,100 μg/ml) (Supplementary Figure 2B, C, D). Accordingly, we detected no amyloid deposition upon incubation of THP-1 macrophages with different concentrations of Scrambled-Aβ42 (Figure 3A, D).

**Figure 3:**
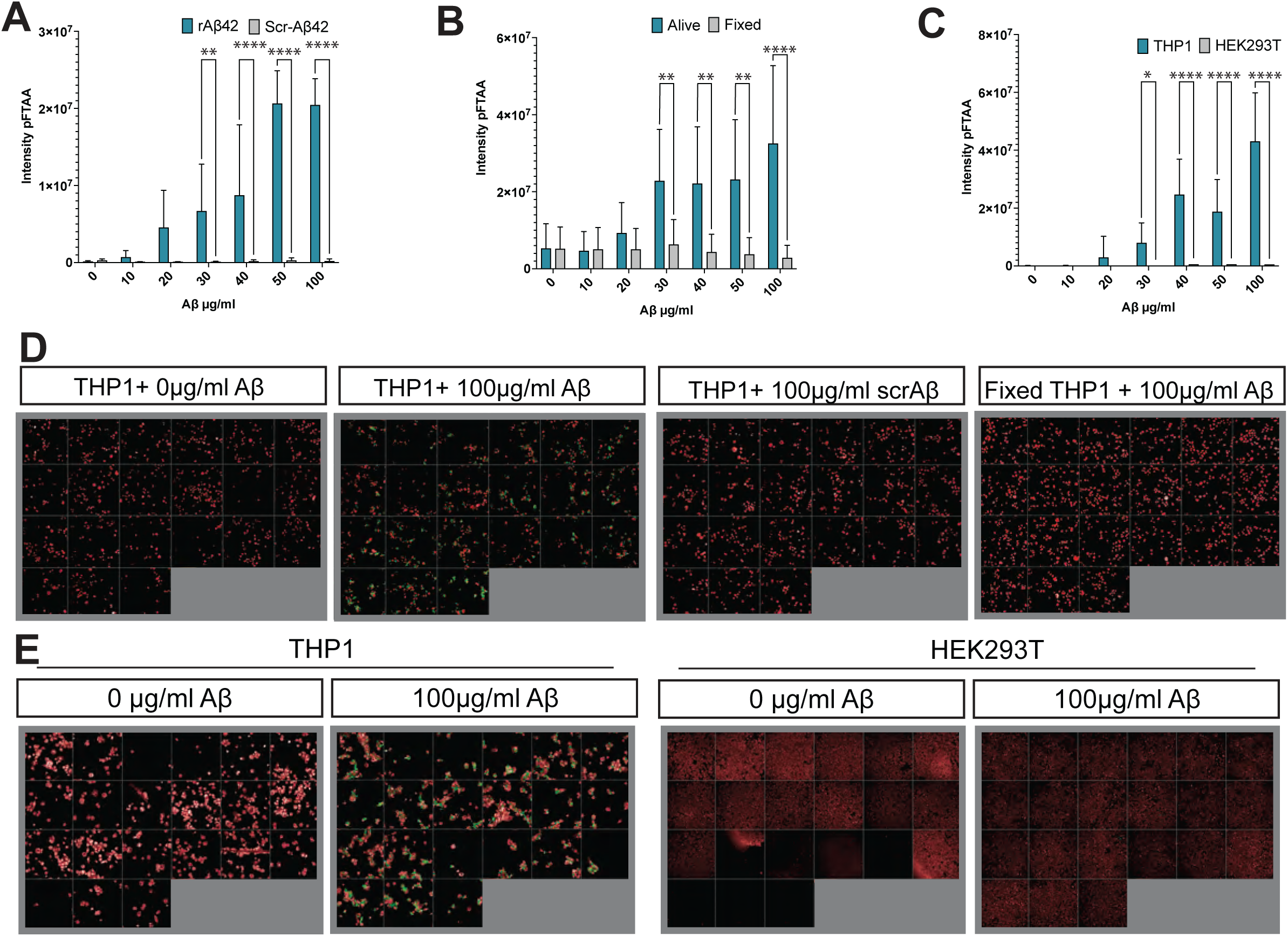
Aggregation propensity and macrophage viability are needed for Aβ42 deposition. A) pFTAA intensity in cells incubated with rAβ42 or scrambled rAβ42. No aggregation is observed in cells treated with scrambled Aβ42. 2 independent experiments with 4 repeats, mean+SD. Statistics: Two-way Anova, with Šídák correction for multiple comparisons. **<0.01, ****<0.0001 B) pFTAA intensity in cells incubated with rAβ42 after or without fixation. 2 independent experiments with 4 repeats, mean+SD. Statistics: Two-way Anova, with Šídák correction for multiple comparisons. **<0.01, ****<0.0001 C) pFTAA intensity of THP-1 macrophages and HEK293T cells treated with increasing concentration of rAβ42. 3 independent experiments with 4 repeats, mean+SD. Statistics: Two-way Anova, with Šídák correction for multiple comparisons. *<0.05, ****<0.0001 D) Representative images of THP-1 cells in different conditions. Each image consists of 21 different fields. Red: nuclei (DAPI), Green: aggregates (pFTAA). E) Representative images of THP-1 and HEK cells incubated with or without 100μg/ml rAβ42. Red: Nuclei (DRAQ7), Green: aggregates (pFTAA)

To ensure that the observed deposition requires cellular activity and is not simply the result of Aβ42 nucleation by cell membranes or other cellular components, we fixed the differentiated THP-1 cells with 4% paraformaldehyde (PFA) before incubation with Aβ42. We chose PFA since it preserves the membrane and does not permeabilize the cells^35^. No deposition was observed in the presence of fixed cells when treated with different concentrations of Aβ42, indicating that cell function plays an essential role in the deposition of Aβ42 in our model (Figure 3B, D). Notably, staining of both live and fixed sAβ42-treated cells with the Aβ42 antibody 6E10 revealed signals under both conditions, indicating the presence of Aβ in distinct aggregating species: monomers in fixed cells and fibrils in living cells (Supplementary Figure 2E). Finally, to ensure that macrophage activity is required for Aβ42 deposition, we tested whether another non-macrophage cell line could cause a similar deposition. We used HEK293T cells for this purpose. When we incubated HEK cells with freshly dissolved Aβ42, we observed some amyloid deposition, but significantly less than that produced by THP-1 cells (Figure 3C, E), consistent with previous results^21^. This indicates that macrophages are particularly prone to this amyloid nucleation phenotype, in line with earlier findings^21^.

Finally, we wanted to confirm these findings using Scanning Electron Microscopy (SEM). Untreated cells (Figure 4 A,B,C) appeared morphologically very distinct from treated cells (Figure 4 D,E,F), again suggesting a cellular response. The space in between cells showed webs of fibrous material in the treated cells (Figure 4F, G, K yellow arrows), that was not found in the sample with untreated cells. Also, amyloid fibrils of Aβ42 grown in a cell-free manner imaged in the same way as the cells, showed a different morphology again (Figure 4I). To ensure that the region with fibril-like assemblies in the treated cells corresponded to the regions of the sample where pFTAA fluorescence was observed, we performed Correlative Light and Electron Microscopy (CLEM, Figure 4J, K, L, M). Cells live stained with pFTAA were first imaged using the light microscope, then after sample preparation the same cells were imaged using SEM. This confirmed that indeed the extracellular fibril-rich web-like structures correspond to regions of high pFTAA fluorescence and can hence be attributed to Aβ42 fibrils generated by the cells.

**Figure 4:**
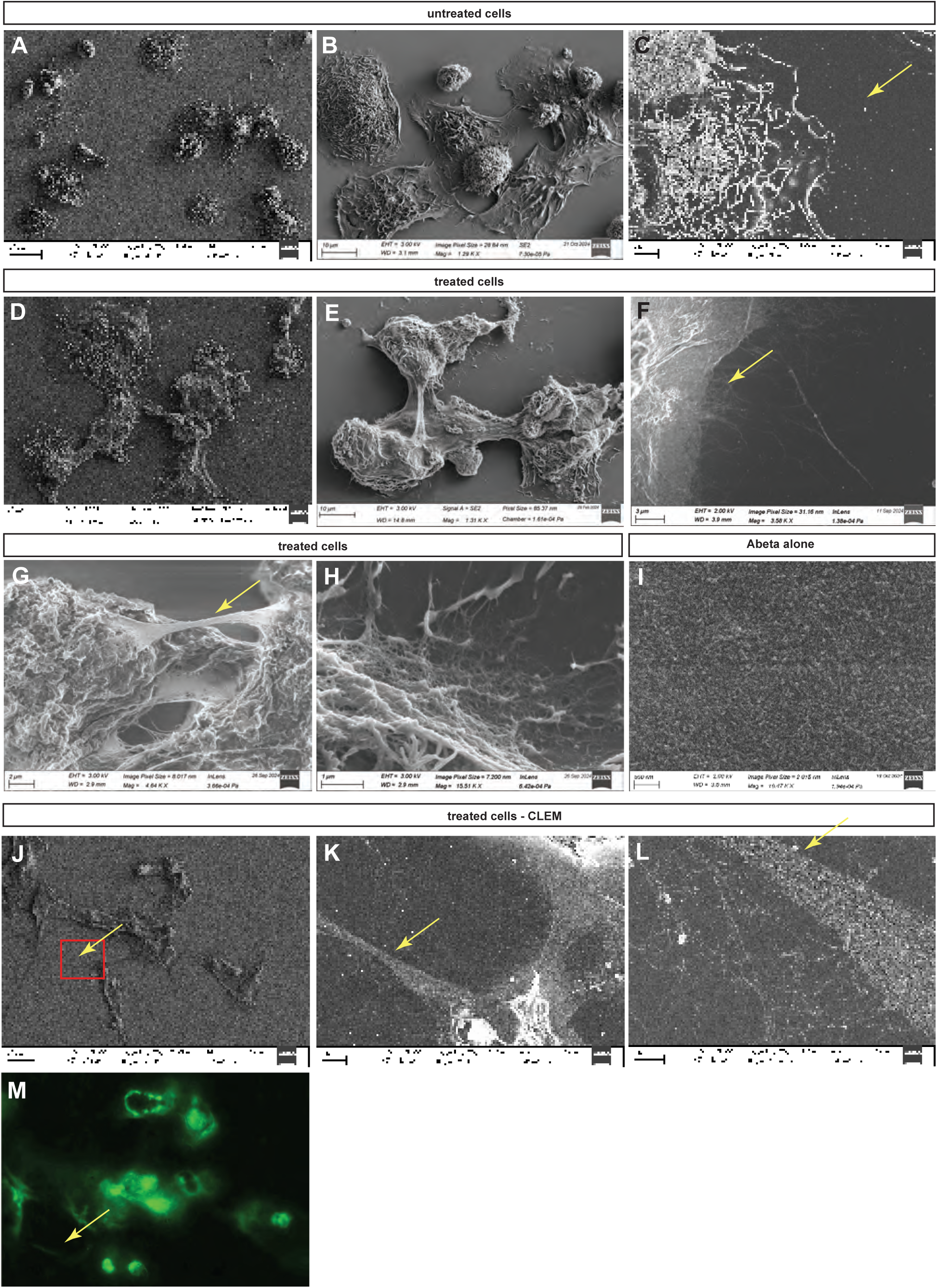
Scanning Electron Microscopy revealed webs of fibril-like material generated by THP-1 cells in response to Aβ42. A-B-C) Untreated THP-1 cells. D-E) Treated THP-1 cells F-G-H) Zoomed-in views of the fibril-like structures formed by THP-1 cells upon sAβ42 treatment. I) Cell-free prepared sAβ42 fibrils imaged in the same manner as the cells. J-K-L-M) Correlative light and electron microscopy combining fluorescence microscopy and SEM of THP-1 cells exposed to sAβ42 and stained with pFTAA.

### Therapeutic antibodies recognize THP-1 amyloid deposition

To determine whether therapeutic antibodies developed to target Aβ fibrils in the human brain could recognize THP-1-generated deposits, we tested four antibodies: Lecanemab, Aducanumab, Donanemab, and Gantenerumab^7,9–11,13,36–38^. For specificity, we compared their binding to THP-1-generated Aβ42 aggregates, in vitro-grown Aβ42 fibrils and seeds, as well as the unrelated Islet Amyloid Polypeptide (IAPP) amyloids, which are associated with Type 2 Diabetes.

Using pFTAA staining, we observed that THP-1-generated Aβ42 aggregates displayed higher fluorescence intensity compared to in vitro-grown fibrils and seeds, suggesting distinct properties and potentially different conformations of cell-generated aggregates (Figure 5A). Staining with IAPP and Aβ antibodies confirmed the deposition of each protein and the specificity of the interaction (Figure 5B, C).

**Figure 5:**
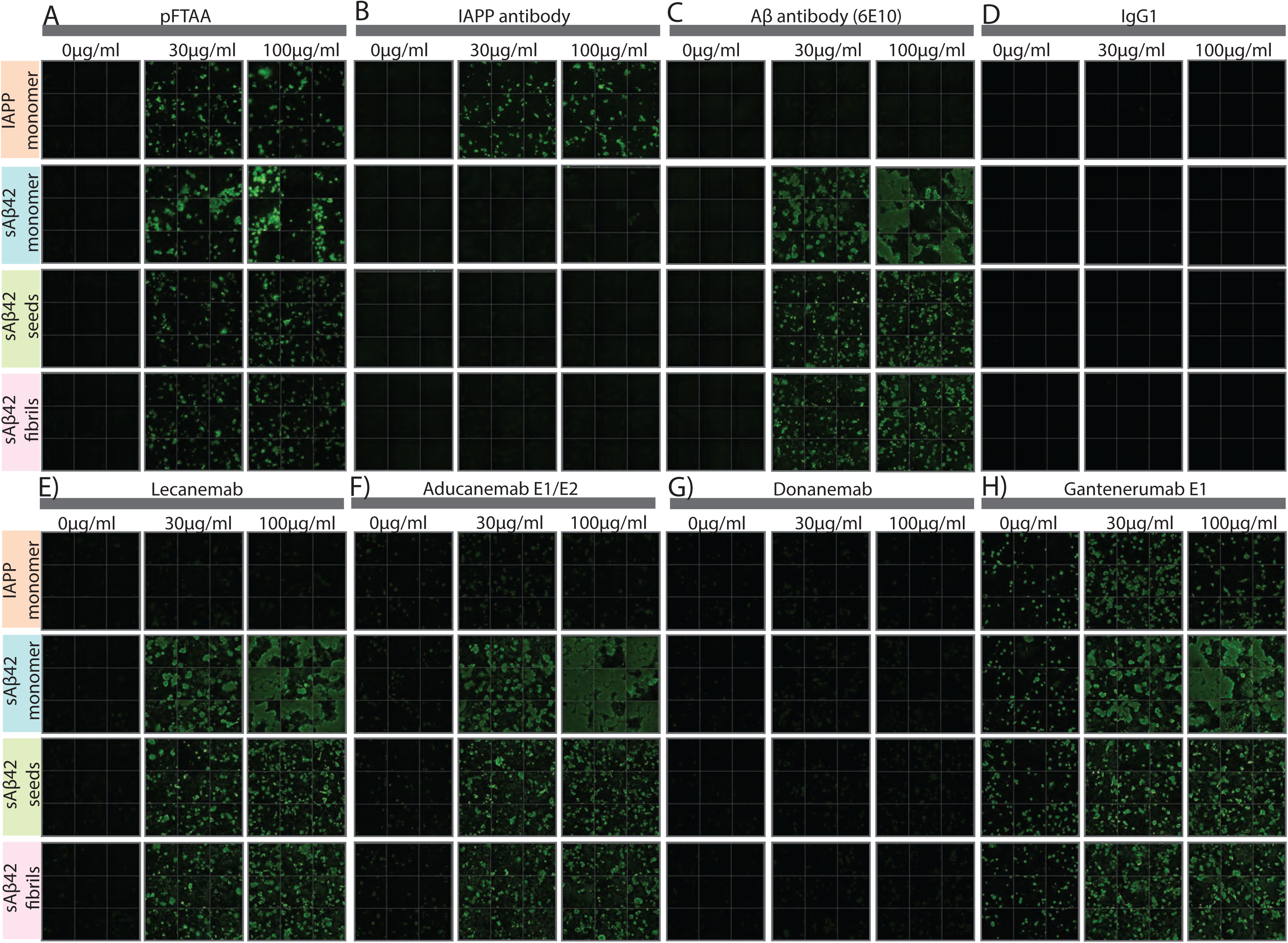
Recognition of Aβ42 aggregates from THP-1 by therapeutic antibodies. A) pFTAA staining of THP-1 incubated with 0,30 and 100μg/ml of IAPP monomer, sAβ42 monomer, sAβ42 cell-free made seeds and fibrils. B) IAPP staining of THP-1 incubated with 0,30 and 100μg/ml of IAPP monomer, sAβ42 monomer, sAβ42 cell-free made seeds and fibrils. C) Aβ staining of THP-1 incubated with 0,30 and 100μg/ml of IAPP monomer, sAβ42 monomer, sAβ42 cell-free made seeds and fibrils. D) Control staining with IgG1 of THP-1 incubated with 0,30 and 100μg/ml of IAPP monomer, sAβ42 monomer, sAβ42 cell-free made seeds and fibrils. E) Lecanemab staining of THP-1 incubated with 0,30 and 100μg/ml of IAPP monomer, sAβ42 monomer, sAβ42 cell-free made seeds and fibrils. F) Aducanemab staining of THP-1 incubated with 0,30 and 100μg/ml of IAPP monomer, sAβ42 monomer, sAβ42 cell-free made seeds and fibrils. G) Donanemab staining of THP-1 incubated with 0,30 and 100μg/ml of IAPP monomer, sAβ42 monomer, sAβ42 cell-free made seeds and fibrils. H) Gantenerumab staining of THP-1 incubated with 0,30 and 100μg/ml of IAPP monomer, sAβ42 monomer, sAβ42 cell-free made seeds and fibrils.

Next, we evaluated antibody binding to THP-1-generated Aβ42 and IAPP aggregates, as well as THP-1 cells incubated with in vitro-grown Aβ fibrils and seeds. Both Lecanemab and Aducanemab specifically bound to Aβ42 aggregates but not to IAPP deposits (Figure 5E, F). Donanemab, which targets pyroglutamate-modified Aβ, did not bind to THP-1-generated Aβ42 aggregates, suggesting that these cells may not produce this specific Aβ variant (Figure 5G). Gantenerumab, however, showed non-specific binding in all conditions, regardless of the presence of Aβ or IAPP (Figure 5H).

### Enhanced seeding and cross-seeding efficiency of THP-1-derived Aβ aggregates

We next evaluated the seeding potential of THP-1cell-derived aggregates. To achieve this, we extracted THP-1 aggregates after 48 hours using a standard fibril extraction protocol^39^ (Figure 6A). These cell-derived aggregates were then compared to cell-free aggregates and patient-derived aggregates in terms of their seeding efficiency. For this comparison, we utilized an Aβ biosensor cell line (HEK293T cells stably expressing mCherry-Aβ42), which produces high fluorescence intracellular spots upon transfection with potent Aβ42 seeds^40^.

**Figure 6:**
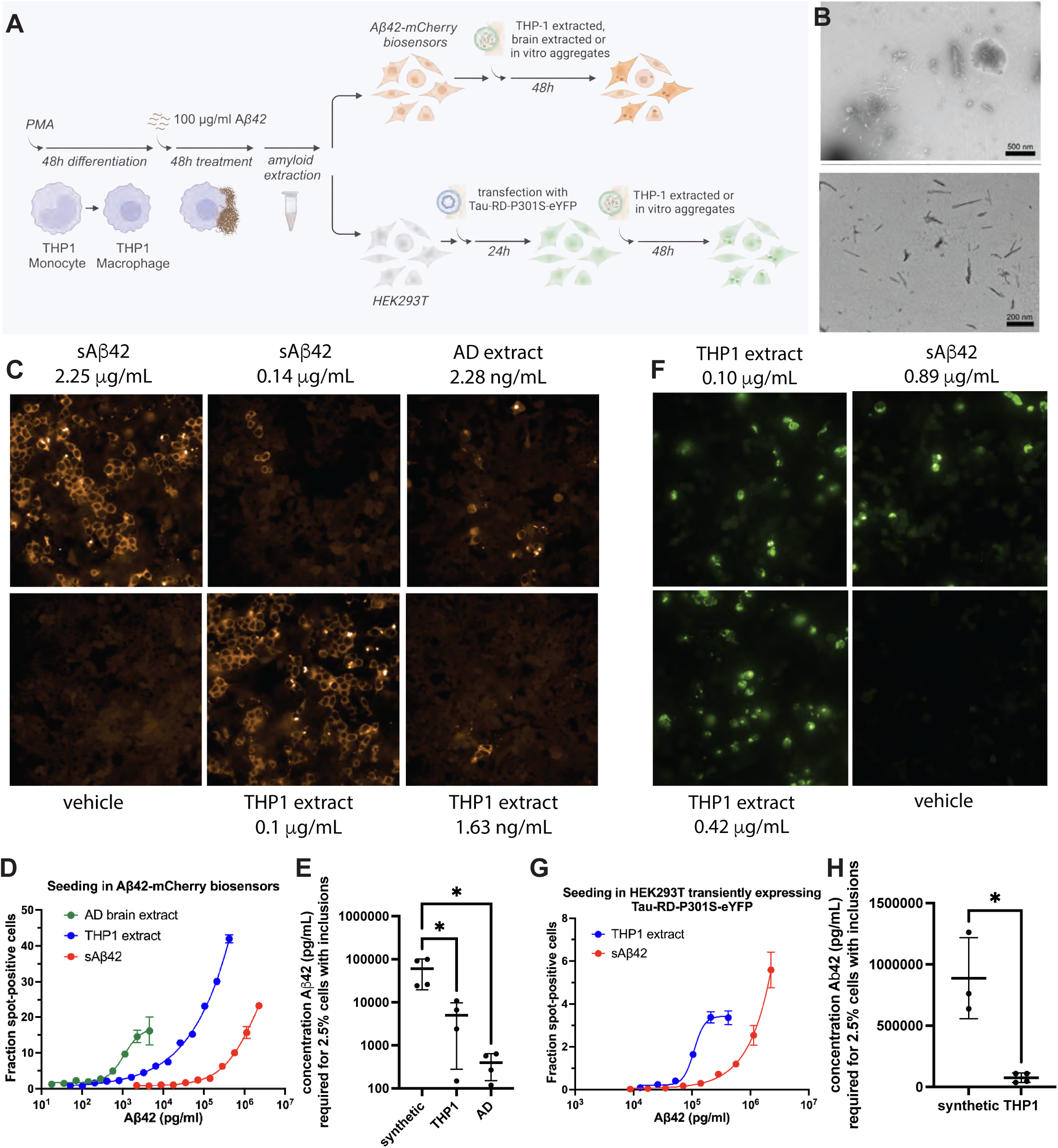
THP-1-derived aggregates have higher seeding activity than cell-free aggregates. A) Schematic of experimental procedures. B) Transmission electron Microscopy of amyloid fragments of Aβ42 extracted from THP-1 cultures. C) Sample fluorescence microscopy images of Aβ biosensor cells treated with human brain extracts from an AD patient brain, amyloid generated in a cell-free manner and amyloids extracted from THP-1 cultures, at the doses indicated. D) Fraction of spot-positive cells in dose-dependent seeding of Aβ42 biosensor cells with human brain extracts from an AD patient brain, amyloid generated in a cell-free manner and amyloids extracted from THP-1 cultures. E) Quantification and statistical comparison of the seeding efficiency of different Aβ preparations in Aβ biosensor cells. Seeding efficiency is expressed at the interpolated concentration of Aβ required to obtain 2.5% aggregate positive cells in the high content assay in D. Comparisons were performed using one way ANOVA followed by pairwise comparison using Dunnett’s test. *** indicates p ≤ 0.001. F) Sample fluorescence microscopy images of HEK293T cells expressing Tau-RD-P301S-eYFP treated with amyloid generated in a cell-free manner and amyloids extracted from THP-1 cultures at the doses indicated. G) Fraction of spot-positive cells in dose-dependent seeding of Tau biosensor cells with the same samples as in C. H) Quantification and statistical comparison of the seeding efficiency in tau biosensor cells, similar to E. Statistical comparison was performed using Welch’s t-test.

The Aβ42 biosensor cells were treated with seeds derived from cell-made fibril extraction, which appeared as fibril fragments by Transmission Electron Microscopy (TEM, Figure 6B). For comparison, we included amyloids extracted from AD patient brain tissue, and synthetic amyloid fibrils prepared in a cell-free manner, by incubating Aβ42 at 1 mg/ml for two weeks at room temperature (Figure 6C). Remarkably, the seeding potential of the THP-1-derived fibrils (Figure 6D) was 12-fold higher than that of the cell-free aggregates (Figure 6D,E), but appeared lower than that of patient-derived material.

We also evaluated the cross-seeding potential of THP-1-derived Aβ42 aggregates and cell-free Aβ42 fibrils on HEK293T cells transiently expressing Tau-RD-P301S-eYFP, the so-called tau biosensor line (Figure 6A). Notably, the THP-1-derived Aβ aggregates exhibited a higher cross-seeding efficiency than cell-free fibrils, although this assay showed more variability than the Aβ biosensor (Figure 6F,G,H). In this biosensor line, we did not compare to patient brain extracts, since these contain tau seeds, precluding conclusions on cross-seeding.

Overall, these findings highlight the significant influence of the cellular environment on the seeding and cross-seeding potential of Aβ42 aggregates. THP-1-derived amyloids demonstrate significantly greater seeding efficiency than cell-free amyloids, yet they still fall short of the potency exhibited by patient-derived aggregates.

### THP-1 and hESC-Derived Microglia Show Similar Early-Stage Responses to Aβ42 Aggregation

We sought to determine whether comparable Aβ deposits could form in the presence of microglia derived from human embryonic stem cells (hESCs). To investigate this, we cultured hESC-derived microglia in the presence and absence of Aβ42 for 48 hours, followed by staining with pFTAA to detect Aβ42 deposition. Our results confirmed the presence of Aβ42 deposits in these cultures as well (Figure 7A). Given that both THP-1 cells and hESC-derived microglia promote Aβ42 aggregation, we next explored whether these two cell types also exhibit similar cellular responses to Aβ42.

**Figure 7:**
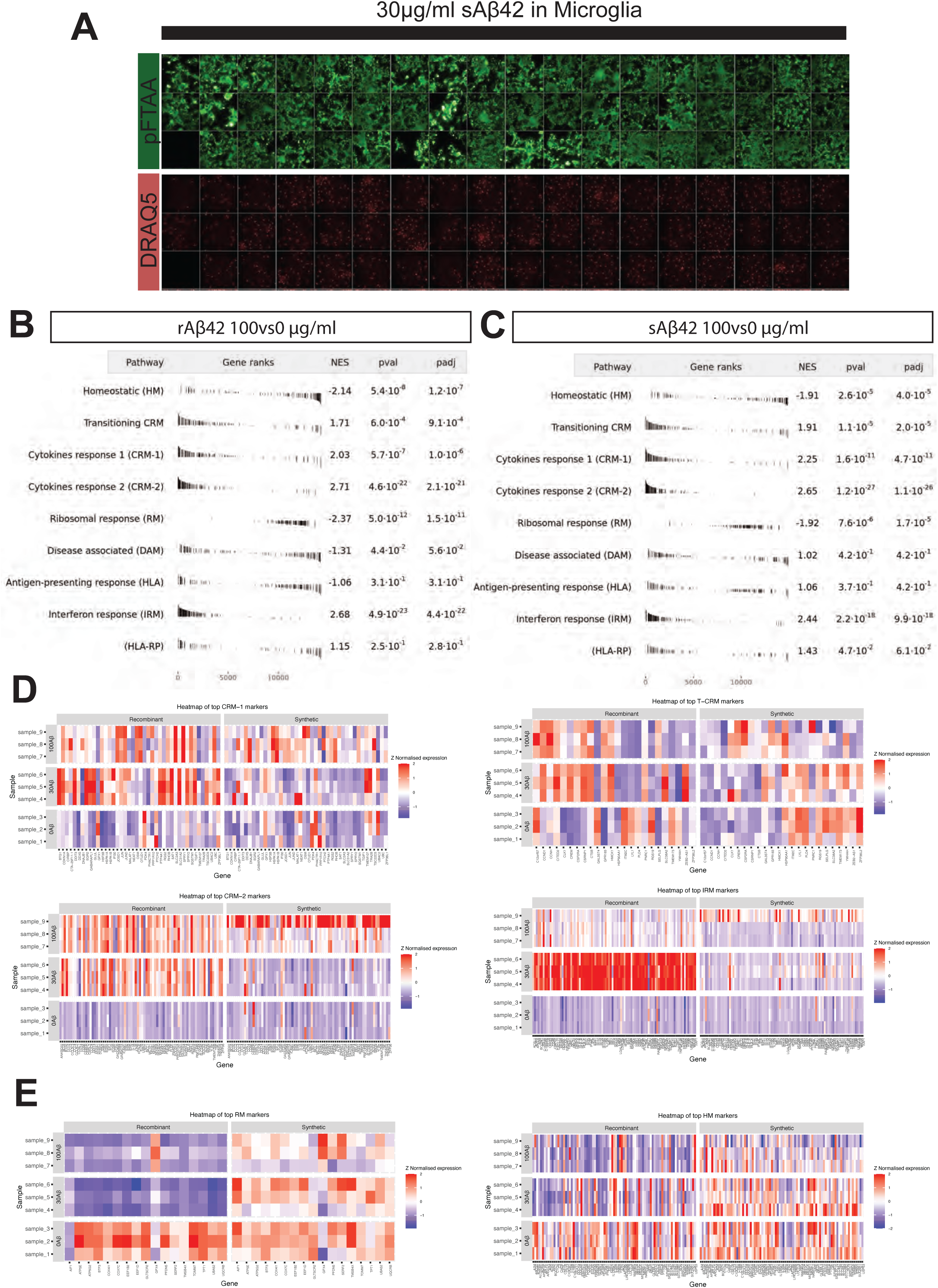
THP-1 response to Aβ42 is similar to microglia in mouse brains. A) pFTAA and DRAQ5 staining of microglia in the presence of 30μg/ml indicating Aβ42 deposition B) GSEA analysis of THP-1 cells treated with rAβ42 (recombinant) and compared to previously identified microglia-Aβ pathways C) GSEA analysis of THP-1 cells treated with sAβ42 (synthetic) and compared to previously identified microglia-Aβ pathways D) Heatmaps of pathways overrepresented in THP-1 cells in response to rAβ42 or sAβ42 E) Heatmaps of pathways underrepresented in THP-1 cells in response to rAβ42 or sAβ42

To assess the cellular response of THP-1 cells to Aβ42 treatment, we conducted transcriptomic analysis on differentiated THP-1 cells exposed to varying concentrations of synthetic (s) or recombinant (r) Aβ42 (0 μg/ml, 30 μg/ml, and 100 μg/ml). This analysis revealed over 1,000 differentially expressed genes, including significant upregulation or downregulation of genes associated with cytokine signaling and inflammatory pathways.

We compared these differentially expressed genes across the two Aβ42 concentrations with established marker genes representing distinct microglial states, as described in the study by Mancuso et al^29^. In their work, human stem cell-derived microglial precursors were xenografted into *AppNL-G-F* mice and characterized for their response to Aβ states after maturation. Our Gene Set Enrichment Analysis (GSEA) revealed a downregulation of genes linked to homeostatic microglia and ribosomal function including P2RY12, CX3CR1 and MAF (Figure 7B, C, E). This occurred alongside upregulation of genes involved in cytokine response pathways (CRM-1 and CRM-2) (Figure 7B, C, D), including upregulation of the proinflammatory cytokines and chemokines IL1B, TNF and CCL3. Notably, we did not observe major alterations in transcriptional signatures associated with Disease-Associated Microglia (DAM) or HLA, potentially because these responses typically emerge in the presence of amyloid plaques during later disease stages, as noted by Mancuso et al ^29,41^. Nevertheless, the transcriptomic changes we identified align with the characteristic microglial responses to oligomeric Aβ forms, indicating that THP-1 cells effectively replicate early-stage disease responses to Aβ42 pathology.

### THP-1 and hESC-Derived Microglia Exhibit Comparable Responses to Genetic Perturbation of TREM2 in Amyloid Deposition

To further validate the THP-1 model as a proxy for microglial responses to amyloid, we assessed the impact of TREM2 perturbation on amyloid deposition using the pFTAA assay. We employed iPSC-derived microglia that carry the Alzheimer’s disease-associated R47H loss-of-function (LOF) mutation in TREM2 and isogenic wild-type lines used earlier^41,42^, alongside THP-1 cells in which TREM2 was knocked out using CRISPR-Cas9 (Figure 8A). In both models, we observed a marked increase in pFTAA-positive amyloid staining compared to their respective controls, indicating enhanced amyloid deposition (Figure 8B,C). This confirms that impaired TREM2 function facilitates the accumulation of extracellular amyloid fibrils also in our cellular system, consistent with its demonstrated role in enhancing AD risk through limiting beneficial microglial functions, including their recruitment to amyloid plaques. These results support the utility of THP-1 cells as a model for probing microglial responses to amyloid, and further indicate that TREM2 plays a conserved role across phagocytic cells in mediating responses to amyloidogenic peptides. This alignment across cell types supports the use of THP-1 cells as a proxy system to explore microglial biology in the context of AD risk genetics.

**Figure 8:**
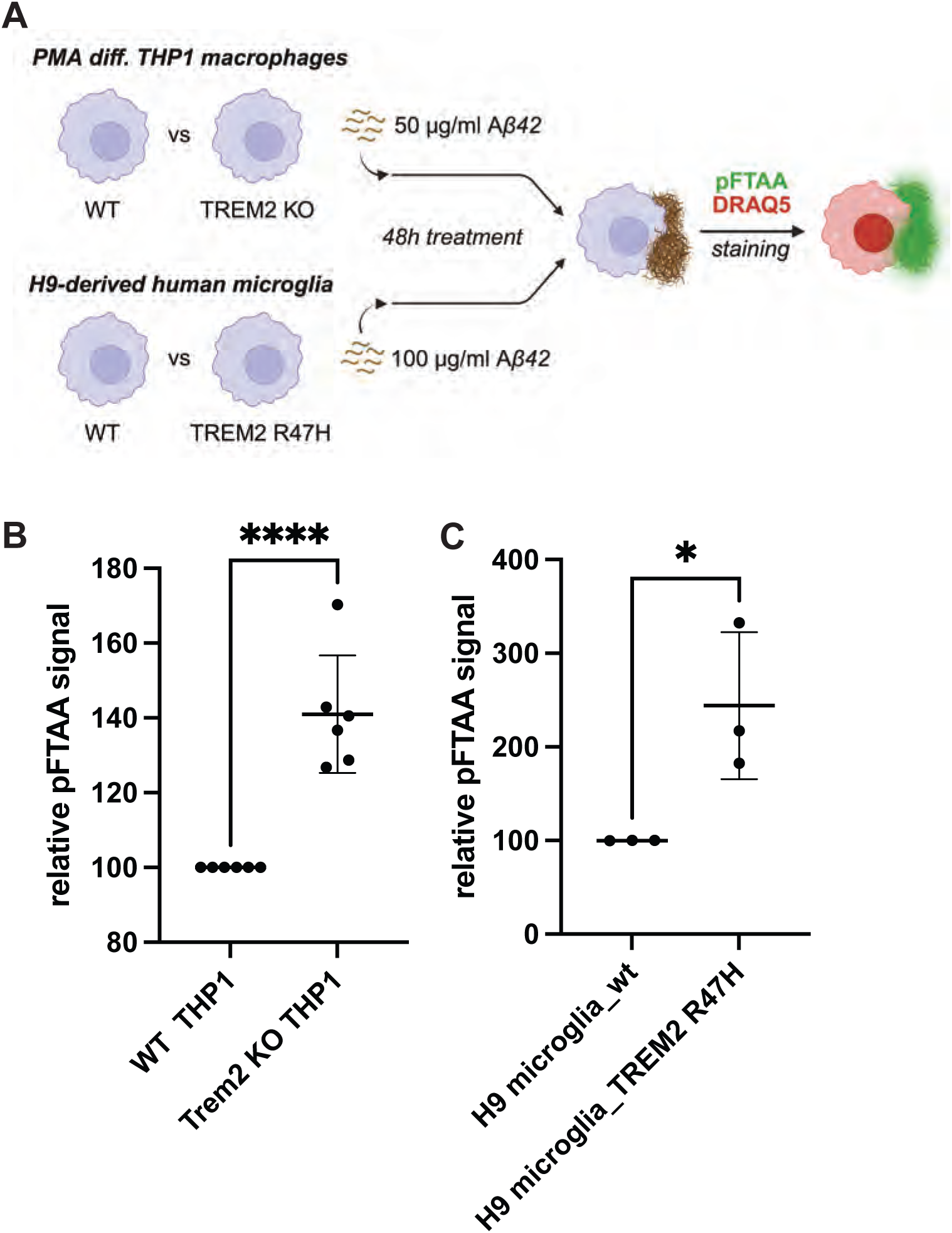
The effect of TREM2 loss of function on in vitro plaque formation by Aβ42 in THP1 cells is similar to hESC-derived microglia. A) Schematic of experimental procedure B) pFTAA staining of THP1 cells treated with 50 μg/mL Aβ42. 6 independent experiments with 5 repeats. Statistics: unpaired t-test C) pFTAA staining of H9-derived human microglia treated with 100 μg/mL Aβ42. 3 independent experiments with 7 repeats. Statistics: unpaired t-test

## Discussion

Our study corroborates recently described observations that microglia contribute to Aβ aggregation^19^. Our data, however, extends these findings to show that macrophage-like cells have a unique capacity in remodelling Aβ into fibrillic structures that are able to seed and propagate amyloid pathology. Our experiments provide a molecular underpinning that microglia, instead of being the main cells involved in clearing Abeta peptides, are in fact generating and initiating the amyloid plaques, outing them strongly upfront in the amyloid cascade. Our work also provides a tractable in vitro model that recapitulates early microglial responses relevant to Alzheimer’s disease (AD) and a way to produce toxic amyloid oligomers in a reproducible way. In particular, we demonstrate that differentiated THP-1 macrophages actively facilitate the extracellular deposition of amyloid fibrils from soluble Aβ42 in a process dependent on cellular activity that remains non-cytotoxic within the experimental timeframe. The resulting cell-derived fibrils exhibit unique structural and functional characteristics, distinct from in vitro cell-free Aβ42 fibrils. Transmission electron microscopy and amyloid dye binding confirmed these structural differences, while functional assays demonstrated enhanced seeding potency for Aβ42 and cross-seeding efficiency for tau. Whether these differences result from THP-1-induced structural polymorphism, associated cofactors, or both remains an open question. Nonetheless, this model effectively recapitulates early microglial responses to Aβ42, as evidenced by transcriptomic profiling, and highlights its relevance for investigating amyloid propagation and inflammatory mechanisms in Alzheimer’s disease (AD). This dual validation—at the levels of functional response and genetic susceptibility—positions THP-1 cells as a valuable model system for early AD biology. Crucially, THP-1 cells not only mirror microglial inflammatory responses but also recapitulate key genetic susceptibility mechanisms. Perturbation of TREM2 function— via the Alzheimer-associated R47H variant in hESC-derived microglia or CRISPR-Cas9 knockout in THP-1 cells—resulted in significantly increased Aβ42 deposition. These results confirm a conserved role for TREM2 across phagocytic cell types in regulating the response to amyloid peptides, and highlight the potential of THP-1 cells for mechanistic dissection of AD risk genes. By capturing both the functional and genetic hallmarks of early microglial-amyloid interactions, the THP-1 model offers a uniquely versatile and scalable platform for preclinical exploration.

These findings build upon earlier studies using Aβ40^21,22^, which is more closely associated with vascular amyloid, but shift the focus to Aβ42, which is more strongly implicated in parenchymal plaque formation and neurotoxicity in AD. While previous research emphasized fibril localization with Aβ40, our study characterizes THP-1-derived Aβ42 aggregates in detail, revealing their potent biological activity. Notably, the fibrils produced by THP-1 cells from Aβ42 demonstrate a 12-fold higher seeding potency compared to cell-free fibrils and a 11-fold increase in tau cross-seeding in biosensor cell lines, underscoring the critical influence of the cellular environment on fibril maturation and pathogenicity. These distinctions emphasize the importance of investigating different Aβ species (or combinations thereof) in cellular systems to better reflect the heterogeneity of amyloid pathology in vivo.

Amyloid deposition required active phagocytic processes, as demonstrated by the lack of fibril formation in fixed THP1 or non-phagocytic HEK293T cells. The extracellular localization of Aβ42 fibrils, confirmed through fluorescence quenching experiments, highlights macrophages’ capacity to promote the accumulation of extracellular aggregates. Our results are in line with the mechanism proposed earlier by Fändrich and co-workers^21^, who suggested that fibril formation is amplified through phagocytic uptake and concentration in endosomal vesicles. However, our data diverge from previous models in showing that extracellular amyloid deposition can occur independently of cell death. This raises a crucial mechanistic question: how do viable phagocytes export amyloid fibrils, and is this process conserved in brain-resident microglia?

Transcriptomic analyses reveal that Aβ42 treatment triggers a robust inflammatory response in THP-1 macrophages, characterized by the upregulation of cytokine signaling and immune activation pathways. This profile closely parallels early microglial activation observed in human microglia xenografted into mouse models exposed to amyloid pathology^29^. The downregulation of homeostatic microglial markers and ribosomal function, alongside the activation of cytokine response pathways, aligns with early-stage responses to oligomeric Aβ^29^. However, the absence of Disease-Associated Microglia (DAM) or antigen presenting transcriptional signatures suggests this model is specific to early disease processes, offering a valuable platform for studying initial inflammatory mechanisms and evaluating therapeutic interventions.

In summary, our study highlights the critical role of macrophage-like cells in driving the extracellular aggregation of Aβ42, revealing key differences between cell-derived and synthetic amyloid fibrils in terms of structure, function, and seeding potential. By effectively mimicking early microglial responses to Aβ42, the THP-1 system offers a uniquely accessible and genetically tractable platform for modeling amyloid-induced inflammation and investigating mechanisms underlying amyloid propagation in AD. The distinct properties of cell-derived fibrils and their recognition by therapeutic antibodies underscore the relevance of this model for preclinical research.

While the THP-1 model offers a tractable and scalable platform to study early amyloid aggregation and inflammatory responses, it has inherent limitations. As a monoculture of peripheral macrophage-like cells, it lacks the full complexity of the CNS microenvironment, including interactions with neurons, astrocytes, and vascular elements that shape microglial behavior in vivo. Moreover, the model does not fully recapitulate the chronic stimulation and tissue remodeling present in neurodegenerative disease. This may explain the absence of later-stage transcriptional states such as disease-associated microglia (DAM) or antigen-presenting phenotypes, which typically emerge in response to dense plaques and long-term pathology. Despite these constraints, the THP-1 model remains uniquely suited for dissecting acute amyloid– microglia interactions and validating genetic risk factors in a human cellular context. Complementary systems—including human iPSC-derived microglia or in vivo models— remain important for modeling the full spectrum of Alzheimer’s disease progression. Future studies using this model could dissect the molecular underpinnings of fibril polymorphism and its impact on amyloid propagation and therapeutic antibody binding. These insights could inform early-stage therapeutic screening and guide the development of conformation-specific treatments for AD.

## Methods

### Cell culture and incubation with Aβ42

THP-1 cells (TIB-202, ATCC) were cultured in RPMI-1640 medium, supplemented with 10% FBS (15517589, FisherScientific), 1mM Sodium Pyruvate (11360070, ThermoFisher Scientific) and 1x non-essential amino acids (12084947, FisherScientific) at 37°C and 5% CO2. Cells were plated at 30000 cells/well in a 96-well plate (PerkinElmer) or in a culture slide (Nunc™ Lab-Tek™ II Chamber Slide™ System) and differentiated with RPMI-1640 supplemented with 50ng/ml phorbol 12-myristate 13-acetate (PMA). After 48h, the medium was changed to RPMI-1640 without PMA. 5h later, the cells were incubated with different concentrations of Aβ42 for 48h. Recombinant Aβ42 (rAβ42, rpeptide) or synthetic Aβ42 (sAβ42, ab120301, Abcam) were prepared by dissolving in sterile H20 and sonicating for 2min. Similarly, Scrambled Aβ42 (rPeptide) was prepared. For the fixed-cells experiments, cells were fixed after the 5-hour rest and before adding Aβ42 with 4% PFA in PBS.

TREM2 knockout THP-1 cell line (and wild type control) are from Abcam (ab269489) and were provided by Jonas Neher. Differentiation and treatment was done as above. HEK293T cells were cultured with DMEM, supplemented with 10% FBS 15517589, FisherScientific), 1mM Sodium Pyruvate (11360070, ThermoFisher Scientific), and 1x non-essential amino acids (12084947, FisherScientific) at 37oC and 5% CO2. Plates were coated with poly-L-lysine, and cells were plated at 30000 cells/well in a 96-well plate (PerkinElmer); 48 h later, the medium was changed for 5h. Cells were then incubated with Aβ42 for 48h.

Human microglia, differentiated from H9-WA09 iPSCs, were treated for 48h with 0,30 and 100μg/ml sAβ42 prepared as described above, with the only difference that half of the TIC medium was changed to contain 2x the sAβ42 concentration.

Aβ peptides used in this study:

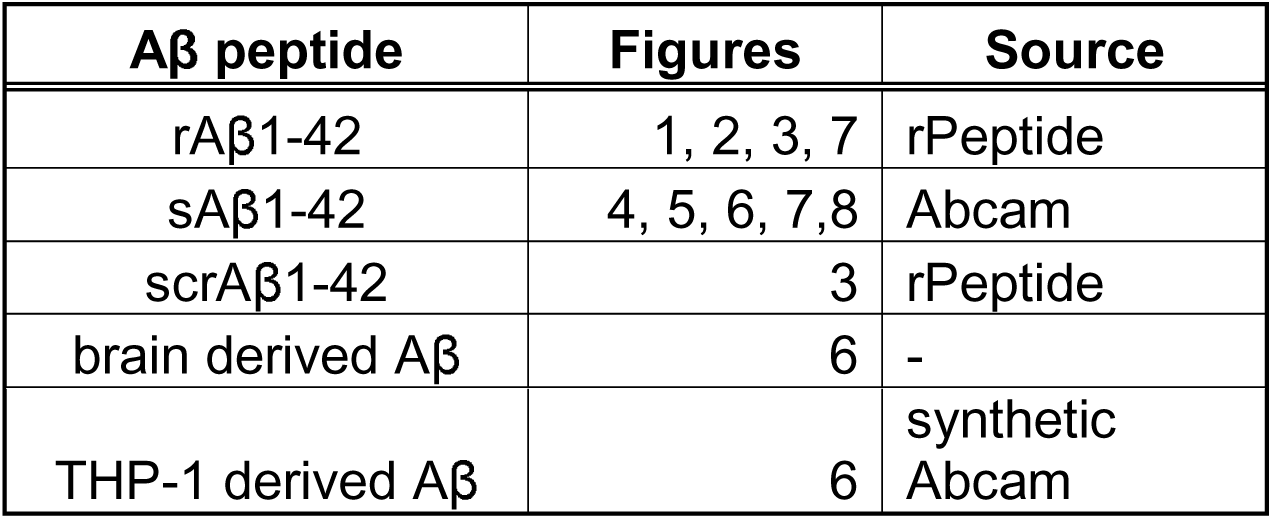

### Congo Red staining

Cells were plated in culture slides (Nunc™ Lab-Tek™ II Chamber Slide™ System) and treated as above. After 48h incubation with Aβ42, cells were washed once with PBS and fixed with ice-cold MeOH for 10 min at 4°C. Cells were incubated for 45min with Congo Red (C6277, Sigma-Aldrich), 0.6%CR (w/v), 80%EtOH (v/v), 3%NaCl (w/v), washed three times with tap water, and incubated with hematoxylin for 6min. Then, washed with 70% EtOH, 3 times with tap water, dehydrated with 90% EtOH, 100% EtOH, two times with 100% Xylol, and mounted. Slides were imaged with Leica DM2500M.

### pFTAA staining

Cells were plated in 96 well plates (PerkinElmer) and treated as above. Cells were washed once with PBS and fixed with 4% formaldehyde for 10min. Cells were stained with 0.5μM pFTAA. Nuclei were stained either with DAPI or DRAQ5 (Biostatus). For live cell staining, cells were incubated with 0.5μM pFTAA and DRAQ5 for 30min at 37°C and 5%CO_2_. Plates were imaged with Operetta CLS: pFTAA (excitation: 390-420nm, emission: 500-550nm), DAPI (excitation: 355-385nm, emission: 430-500nm) or DRAQ5 (excitation: 615-645nm, emission: 655-760nm) using 21 fields and 3 stacks (−2 to 2μm) and analyzed with Harmony software. Nuclei were found using the DAPI or DRAQ5 channel. The threshold for identifying pFTAA-positive image regions was set at an intensity of 3000 and above.

### Antibody staining

Cells were plated in 96 well plates (PerkinElmer) and treated as above. Cells were washed once with PBS and fixed with 4% formaldehyde for 10mn. Cells were blocked for 1h with 10% goat serum in PBS with 0.05% Tween20. Then, cells were incubated with 10ug/ml primary antibodies for 1 h in blocking buffer at room temperature. Cells were washed three times with PBS with 0.05% Tween 20 and incubated for 1h with 1:500 goat anti-Human-Alexa 488 (Invitrogen, A-101013) antibody at room temperature in blocking buffer. Finally, they were stained for 30 minutes with DRAQ5 (Biostatus).

### LDH release

LDH release was quantified using CyQUANT™ LDH Cytotoxicity Assay, according to the manufacturer. Absorbance was measured at 490 and 680nm using an Omega plate reader (BMG Labtech). To determine the LDH release, 680nm absorbance was subtracted from 490nm. Values were then normalized based on lysed cells (100% LDH release).

### Trypan Blue quenching

Live cells were imaged with Operetta CLS or Nikon TiE A1R. For Operetta CLS, live imaging temperature was set at 37°C and environment CO_2_ at 5%. After imaging pFTAA staining, 30μl of Trypan Blue (Biorad) was added, and wells were imaged again. Imaging of the wells after the addition of Trypan blue was concluded within 5 min. Confocal imaging was done at 37 ^°^C and 5% CO_2_ with an Eclipse Ti confocal with Plan APO VC 10× DIC N1 lens, with excitation using a 487.8 nm laser line, and emission was collected at 500–550 nm.

Transfection of 30,000 HEK293T cells per well were plated in 96-well plates (PerkinElmer) and were transfected with 100ng GFP plasmid (Lonza) using Lipofectamine 3000 (Invitrogen) according to the manufacturer’s instructions. After 48h, the medium was changed, and cells were imaged live in the presence and absence of Trypan blue, as discussed above.

### Thioflavin T fluorescence

rAβ42 and Scrambled-Aβ42 were prepared in different concentrations in RPMI1640+ 10%FBS with 25μM ThT. Samples were pipetted in μclear medium binding half area plates (Greiner, #675096) and measured over time with Fluostar (BMG Labtech) at 37°C at excitation 440nm and emission at 480nm.

### Transmission Electron Microscopy

Samples resulting from ThT kinetics were spotted in a copper grid (Formvar/Carbon on 400 Mesh Copper-Agar Sci, AGS162-4), previously glow discharged for 30 sec, and left to be absorbed for 3 min. After this, the grid was washed 3 times with MQ and negatively stained with an aqueous solution of uranyl acetate (2%) in aqueous solution for 1min. The grid was washed with MQ and imaged with a transmission electron microscope (JEM-1400 Jeol, Japan), operated at 80kV, equipped with an Olympus SIS Quemesa 11 MP camera.

### Scanning Electron Microscopy (SEM)

Cells cultured on coverslips as indicated above were fixed with 4% paraformaldehyde (PFA)+ 2.5% glutaraldehyde (GA) in 0,1M sodium cacodylate buffer (pH 7.2) or 0,1 M HEPES buffer (pH 7.4) first for one hour by adding double concentrated fixative in 1:1 proportion to the culture medium, followed by overnight fixation with 4% PFA+ 2.5% GA in 0,1M buffer at 4°C. The next day, the coverslips were rinsed with 0,1 M buffer, 3 x 5 minutes and subsequently postfixed with 1% osmium tetroxide in 0,1M buffer for 1 hour on ice. Next, they were washed 5 x 3 minutes with milliQ water and dehydrated with graded series of ethanol: 1 × 10 minutes each step (30, 50, 70, 90, 100, 100 % ethanol) at RT followed by critical point drying (Leica CPD300, Wetzlar, Germany) in pure ethanol. The coverslips were mounted on the SEM stubs with carbon tape and sputter coated with 5nm chromium followed by 5nm of carbon (Leica ACE600, Wetzlar, Germany).

The samples were imaged with Scanning Electron Microscope (Zeiss Sigma VP; Oberkochen, Germany) operated at 2-3 kV, using secondary electron detector or InLens detector.

### Light Microscopy Imaging of pFTAA fluorescence

For image acquisition, a Nikon NiE upright microscope was used equipped with a Yokogawa CSU-X spinning-disk module with Prime BSI camera (Teledyne Photometrics) in combination with a 20x Fluor water dipping objective (NA 0,5). The setup was controlled by NIS-Elements (5.42.04, Nikon Instruments Europe B.V.).

Green fluorescence of pFTAA was excited with Blue diode (488, sapphire 50mW) and collected with 525/50 filter.

For post processing, NIS-Elements (6.10.01, Nikon Instruments Europe B.V.) was used. The images were clarified and denoised using clarify.ai and denoise.ai.

### Aggregation Prediction

APR and aggregation tendency for Aβ42 and scrΑβ42 were performed using TANGO, an algorithm for predicting aggregation-prone regions, using the sequences provided by rPeptide.

### Fibril extraction from THP1 cells

Cells were collected in Tris-EDTA buffer. They were homogenized using FastPrep (MP Biosciences) following centrifugation for 5’ at 3100g and 4°C. The resulting pellet was resuspended in Tris-EDTA and the above process was repeated 9 times. The pellet was then washed 9 times with H20, using FastPrep homogenization following centrifugation (5’ at 3100g and 4°C). The supernatant was recovered after every wash.

### Extraction of tau aggregates

Following approval, brain tissue from autopsy cases was received from UZ/KU Leuven Biobank. The tissue samples were all fresh-frozen samples of frontal cortex. Immunohistochemical evaluation of formalin-fixed, paraffin-embedded sections from the same anatomical region (i.e. frontal cortex) confirmed the presence of abundant tau-immunoreactive pathology with characteristic features of each condition. Sarkosyl-insoluble material was extracted from cortex tissue as described in previous work^43,44^. Briefly, tissue homogenisation was performed with a FastPrep (MP Biomedicals) in 10 volumes (w/v) cold buffer (10 mM Tris-HCl pH 7.4, 0.8 M NaCl, 1 mM EGTA and 10% sucrose) and a centrifugation step at 20,000 x g for 20 min at 4°C. Universal Nuclease (Pierce) was added to the supernatant, followed by a 30 min incubation at room temperature. Subsequently, the sample was brought to 1% Sarkosyl (Sigma) and incubated for 1h at room temperature while shaking (400 rpm), followed by centrifugation at 350,000 x g for 1h at 4°C. The pellet was washed once, resuspended in 50 mM Tris-HCl pH 7.4 (175 mg of starting material per 100 µl) and stored at −80°C.

### Aβ42 biosensor seeding assay

We used the biosensor line previously described^40^. On day 1, mCherry Aβ42 HEK biosensor cells were plated in a poly-L-lysine-coated CellCarrier 96 Ultra Plate at a density of 12,000 cells/well. After 24h, samples (THP-1 extracted aggregates, brain extracted aggregates, cell-free made aggregates) were sonicated using a Bioruptor Pico, and cells were transfected with Lipofectamine 3000 mixed with seeds or buffer. After 48h, cells were fixed with 4% formaldehyde and stained with DAPI before imaging with the Operetta system in DAPI, mCherry, and DPC channels. For phenotype analysis, an intensity cutoff of >2000 was used to include both cells with higher mCherry intensity and those with spots. Aβ42 concentrations were measured using the Amyloid Beta 42 Human ELISA Kit (Invitrogen KHB3441, Fisher 10602644), with samples prepared through a series of dilutions.

### Tau biosensor seeding assay

On day 1, HEK29ET cells were plated in a poly-L-lysine-coated CellCarrier 96 Ultra Plate at a density of 10,000 cells/well. After 24h, cells were transfected using Lipofectamine3000 with TauRD-P301S-eYFP. After 24h, samples (THP-1 extracted aggregates, brain extracted aggregates, cell-free made aggregates) were sonicated using a Bioruptor Pico, and cells were transfected with Lipofectamine 3000 mixed with seeds or buffer. After 48h, cells were fixed with methanol and stained with DAPI before imaging with the Operetta system in DAPI, YFP and DPC channel.

### Preparation of Aducanumab, Lecanemab, Gantenerumab and Donanemab

The variable domain amino acid sequences for these antibodies were obtained from the KEGG DRUG Database. Synthetic genes encoding the respective variable domains were synthesized by Twist Biosciences (South San Francisco, CA, USA) and cloned into the pTRIOZ-hIgG vector (Invivogen, San Diego, CA, USA). The VL domain was inserted using AscI/BsiWI restriction sites, while the VH domain was inserted using AgeI/NheI. The open reading frames of the cloned antibodies were verified through Sanger sequencing (LGC Genomics, Berlin, Germany). CHO cells (A29127, Thermo Fisher Scientific) were transiently transfected with the plasmid DNA using the TransIT-PRO® Transfection Reagent (MIR 5740, Mirus Bio), following the manufacturer’s instructions. The transfected cells were cultured in suspension under agitation at 32°C for 14 days. Supernatants were collected two weeks post-transfection and incubated overnight at 4°C with AmMag™ Protein A Magnetic Beads (L00939, GenScript). Antibodies were purified using the AmMag™ SA Plus system (L01013, GenScript) after magnetic bead separation. The purity of the antibodies was assessed by densitometric analysis of Coomassie Blue-stained SDS-PAGE gels under non-reducing conditions, with all antibodies showing an estimated purity above 75%.

### RNA sequencing

#### RNA quality control

For samples THP1_0Ab (1), THP1_0Ab (2), THP1_0Ab (3), THP1_30Ab (1), THP1_30Ab (2), THP1_30Ab (3), THP1_100Ab (1), THP1_100Ab (2), THP1_100Ab (3), RNA concentration and purity were determined spectrophotometrically using the Nanodrop ND-8000 (Nanodrop Technologies) and RNA integrity and concentration were assessed using a Fragment Analyzer SS-RNA kit (Agilent). For samples THP1_0sAb (1), THP1_0sAb (2), THP1_0sAb (3), THP1_30sAb (1), THP1_30sAb (2), THP1_30sAb (3), THP1_100sAb (1), THP1_100sAb (2), and THP1_100sAb (3), RNA concentration and purity were determined spectrophotometrically using the Nanodrop ND-8000 (Nanodrop Technologies) and RNA integrity and concentration were assessed using a BioAnalyzer Total RNA Eukaryote Nano assay (Agilent).

#### Library Preparation

Per sample, an amount of 500 ng of total RNA was used as input. Using the Illumina Stranded mRNA Sample Prep Kit (protocol version: # 1000000124518 v02 (April 2021)) poly-A containing mRNA molecules were purified from the total RNA input using poly-dT oligo-attached magnetic beads. The purified mRNA was fragmented and in a reverse transcription reaction using random primers and Actinomycin D, RNA was converted into first strand cDNA and subsequently converted into double-stranded cDNA in a second strand cDNA synthesis reaction using dUTP to achieve strand specificity. The cDNA fragments were extended with a single ‘A’ base to the 3’ ends of the blunt-ended cDNA fragments after which pre-index anchors were ligated preparing the fragments for dual indexing. Anchor-ligated fragments were then purified using magnetic beads. Finally, enrichment PCR was carried out to enrich those DNA fragments that have anchor-ligated DNA fragments and to add indexes and primer sequences for cluster generation.

#### Sequencing

Purified dual-indexed sequence-libraries of each sample were equimolarly pooled, converted with the Adept PCR Plus kit (Element Biosciences) and sequenced on Element Biosciences AVITI (two runs, 2×75 Cloudbreak High kit, single read 100 (101-10-10-0)) at the VIB Nucleomics Core (https://nucleomicscore.sites.vib.be/en).

### Data analysis

#### Preprocessing

Low quality ends and adapter sequences were trimmed off from the sequenced reads with Cutadapt 3.2 ^45,46^. Subsequently, small reads (length < 35 bp), polyA-reads (more than 90 % of the bases equal A), ambiguous reads (containing N), low-quality reads (more than 50 % of the bases < Q25) and artifact reads (all but three bases in the read equal one base type) were filtered using using FastX 0.0.14 and ShortRead 1.58.0 ^47^. With Bowtie2 2.4.5 we identified and removed reads that align to phix_illumina^48^.

#### Mapping

The preprocessed reads were aligned with STAR aligner v2.5.2b to the reference genome of Homo sapiens (GRCh38.88)^49^. Default STAR aligner parameter settings were used, except for ‘--outSAMprimaryFlag OneBestScore --twopassMode Basic --alignIntronMin 50 –alignIntronMax 500000 --outSAMtype BAM SortedByCoordinate’. Using Samtools 1.15.1, reads with a mapping quality smaller than 20 were removed from the alignments^50^.

#### Counting

The number of reads in the alignments that overlap with gene features were counted with featureCounts 1.5.3^51^. Following parameters were chosen: -Q 0 -s 2 -t exon -g gene_id. We removed genes for which all samples had less than 1 count-per-million. Raw counts were further corrected within samples for GC-content and between samples using full quantile normalization, as implemented in the EDASeq package from Bioconductor^52^.

#### Differential gene expression

Using the EdgeR 3.42.4 package of Bioconductor, a negative binomial generalized linear model (GLM) was fitted against the normalized counts^53^. We did not use the normalized counts directly, but worked with offsets. Differential expression was tested for with a GLM likelihood ratio test, also implemented in the EdgeR package. The resulting p-values were corrected for multiple testing with Benjamini-Hochberg to control the false discovery rate^54^.

### Fast Gene Set Enrichment Analysis

Gene set analyses were performed using the R implementation of the Fast Gene Set Enrichment Analysis algorithm^55^. Microglial state markers were defined as the top one-hundred strongest differentially expressed genes by p-value for each microglial state as quantified and reported by Mancuso et al.^29^ Differentially expressed genes from Thp1 cells were then compared to these states by their log2 fold change using the fgsea function.

### Embryonic Stem Cell Culture

The experiments were performed using the established H9 stem cell line (WAe009-A; https://hpscreg.eu/cellline/WAe009-A). Routine culturing and maintenance of the stem cells were performed using the Cellartis DEF-CS 500 Culture System (Takara; #Y30010 & #Y30017). Cells were maintained in a humidified chamber at 37°C with 5% CO.

### Microglial differentiation and maturation

Human microglial-like cells were generated using the protocol of Fattorelli *et al.* 2021^56^. H9-TREM2^R47H^ were developed previously ^42^. In short, on day 0, H9-WA09 or H9-TREM2^R47H^ hESCs were plated at a density of 46,000 cells/cm² in a U-bottom low adherent 96-well plate (Sigma, #CLS7007) in BVS embryonic body (EB) medium, consisting of mTeSR1 (Stemcell, #85850), 50 ng/ml BMP4 (Peprotech, #120-05), 50 ng/ml VEGF (Peprotech, #100-20), and 20 ng/ml SCF (Peprotech, #300-03). Over a period of 3 days, EBs were formed with the medium changed daily.

On day 4, the EBs were carefully collected in a 50 ml Falcon tube and transferred to a 6-well plate at a density of roughly 20 EBs per well. The medium was then changed to SMIFT differentiation medium, which includes X-VIVO 15 (Lonza, #LO BE02-060Q), 2 mM GlutaMAX (Thermo Fisher, #35050038), 100 U/ml Pen/Strep (Sigma, #P4333), 50 µM 2-mercaptoethanol (Thermo Fisher, #31350010), 50 ng/ml SCF (Peprotech, #300-03), 50 ng/ml M-CSF (Peprotech, #300-25), 50 ng/ml IL-3 (Peprotech, #200-03), 50 ng/ml FLT3 (Peprotech, #300-19), and 5 ng/ml TPO (Peprotech, #300-18). The medium was changed after 4 days.

During the medium change on day 8, the free-floating EBs were returned to their wells. On day 11, the medium was changed to FGM differentiation medium, which contains X-VIVO 15 (Lonza, #LO BE02-060Q), 2 mM GlutaMAX (Thermo Fisher, #35050038), 100 U/ml Pen/Strep (Sigma, #P4333), 50 µM 2-mercaptoethanol (Thermo Fisher, #31350010), 50 ng/ml FLT3 (Peprotech, #300-19), 50 ng/ml M-CSF (Peprotech, #300-25), and 25 ng/ml GM-CSF (Peprotech, #300-03).

On day 18, the first batch of microglial progenitors was collected using a 37 µM reversible strainer (Stemcell, #27250). This strainer allows the EBs to be replated in FGM differentiation medium, enabling the collection of two additional batches of microglial progenitors, each 7 days apart.

For further microglial maturation, the microglial progenitors were spun down at 300g for 5 minutes and plated at a density of 20,000 cells/cm² in a PLO/laminin-coated 96-well plate (Thermo Fisher, #165305) in TIC medium. The TIC medium composition is as follows: DMEM/F12 (Thermo Fisher, #11330057), 100 U/ml Pen/Strep (Sigma, #P4333), 200 mM L-glutamine (Sigma, #G7513), 5 µg/ml Insulin (Sigma, #I9278), 5 ng/ml N-Acetyl-L-cysteine (Sigma, #A7250), 50 mg/ml apo-transferrin (Sigma, #T1147), 20 µg/ml sodium selenite (Sigma, #S9133), 1 µg/ml heparin sulfate (Amsbio, #GAG-HS01), 50 ng/ml M-CSF (Peprotech, #300-25), 100 ng/ml IL-34 (Peprotech, #200-34), 2 ng/ml TGF-β (Peprotech, #100-21), 10 ng/ml CX3CL1 (Peprotech, #300-31), and 1.5 ng/ml cholesterol (Sigma, #C4951). The medium was changed after 3 days, and experiments/stimulations were performed after 5 days after plating.

### Statistics

Statistical analysis was done using GraphPad Prism versions 9 and 10. The statistical tests and significance levels are indicated in the figure legends.

## Ethics approval and consent to participate

Ethical approval to access and work on the human tissue samples was given by the UZ Leuven ethical committee (Leuven/Belgium; File-No. S63759). An informed consent for autopsy and scientific use of autopsy tissue with clinical information was granted from all subjects involved (case A-21270 with LC ID 93).

## Availability of data and materials

All data generated or analysed during this study are included in this published article and its supplementary information files.

## Authors’ contributions

KK, MR, BDS, FR and JS conceived the study. KK, MR, SB, DT, MS, GA performed experiments. DM, BH, MF, MR, KK, JS, FR performed analyses. BP, MDW, DRT, MW, JN contributed materials and samples. KK, FR, JS wrote the manuscript and all authors provided feedback.

## Acknowledgments

The authors gratefully acknowledge the Bio Imaging Core, Department of Neurosciences KU Leuven, VIB – KU Leuven Center for Brain & Disease Research for their support and assistance in this work. Light and electron microscopy imaging was performed at the VIB Bioimaging Core Leuven. In particular, the authors thank Nikky Corthout of the VIB Bio Imaging Core Leuven for her support & assistance with light microscopy imaging. Library preparation, sequencing and statistical data analysis were performed by VIB Nucleomics Core (https://nucleomicscore.sites.vib.be/en).

