## supplementary materials for "Phagocytes as Plaque Catalysts: Human Macrophages Actively Generate Pathogenic Aβ42 Fibrils with Seeding and Cross-Seeding Potency"

### Supplementary Figures

#### Supplementary Figure 1

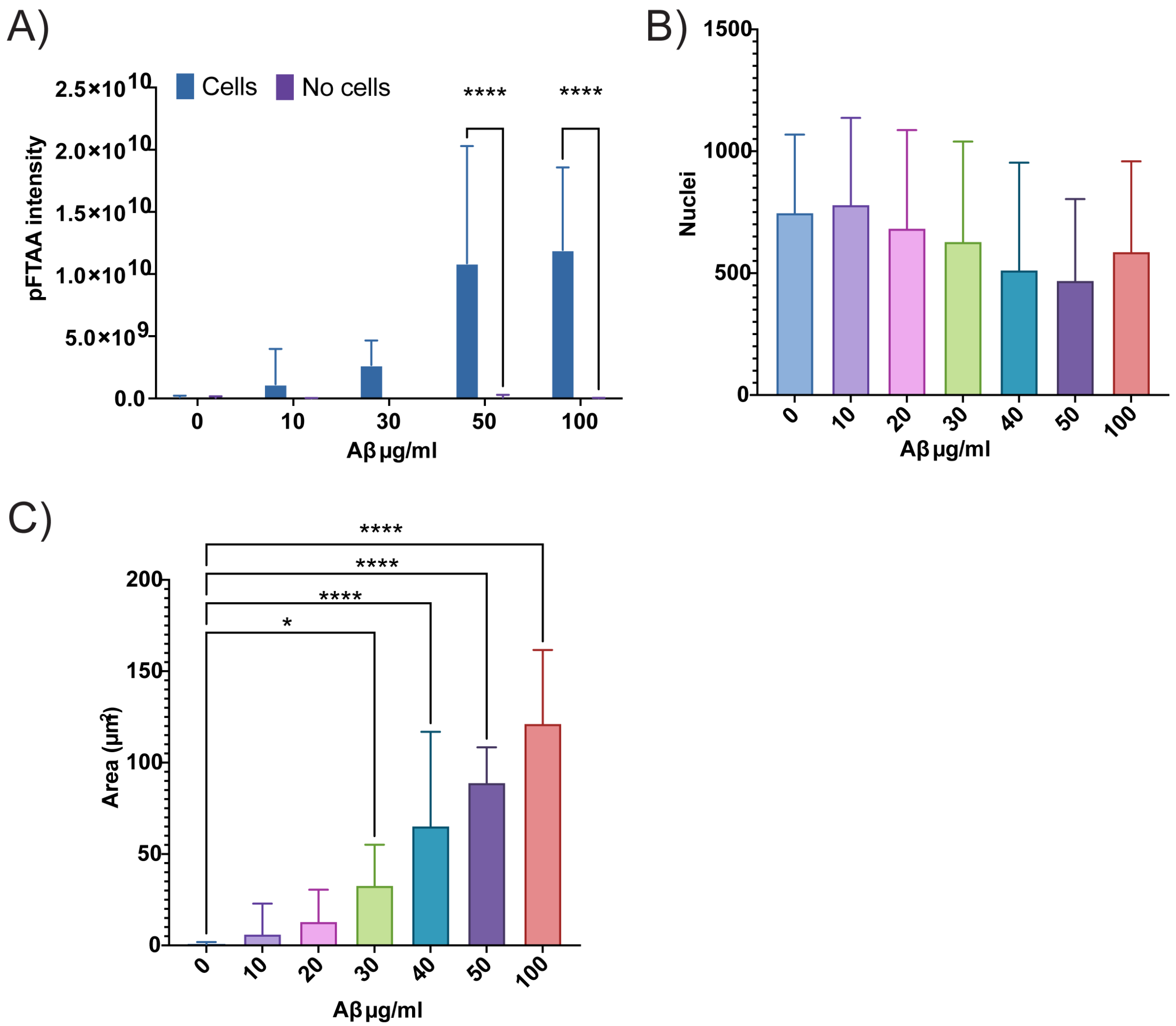


**Supplementary figure 1. Deposition of Aβ in presence or absense of cells**
A) Total intensity of pFTAA per condition in presence and absence of cells. 3 independent experiments with 4 repeats. Mean+SD. Statistics: Two-way Anova with Šídák correction for multiple comparisons. ****<0.0001
B) Number of cells. 4 independent experiments with 4 repeats, mean+SD. Statistics: Ordinary one-way ANOVA with Dunnett correction for multiple comparisons.
C) pFTAA stained area in wells of THP1 cells incubated with different concentrations of Aβ. 4 independent experiments with 4 repeats, mean+SD. Statistics: Ordinary one-way ANOVA with Dunnett correction for multiple comparisons. *<0.05, ****<0.0001

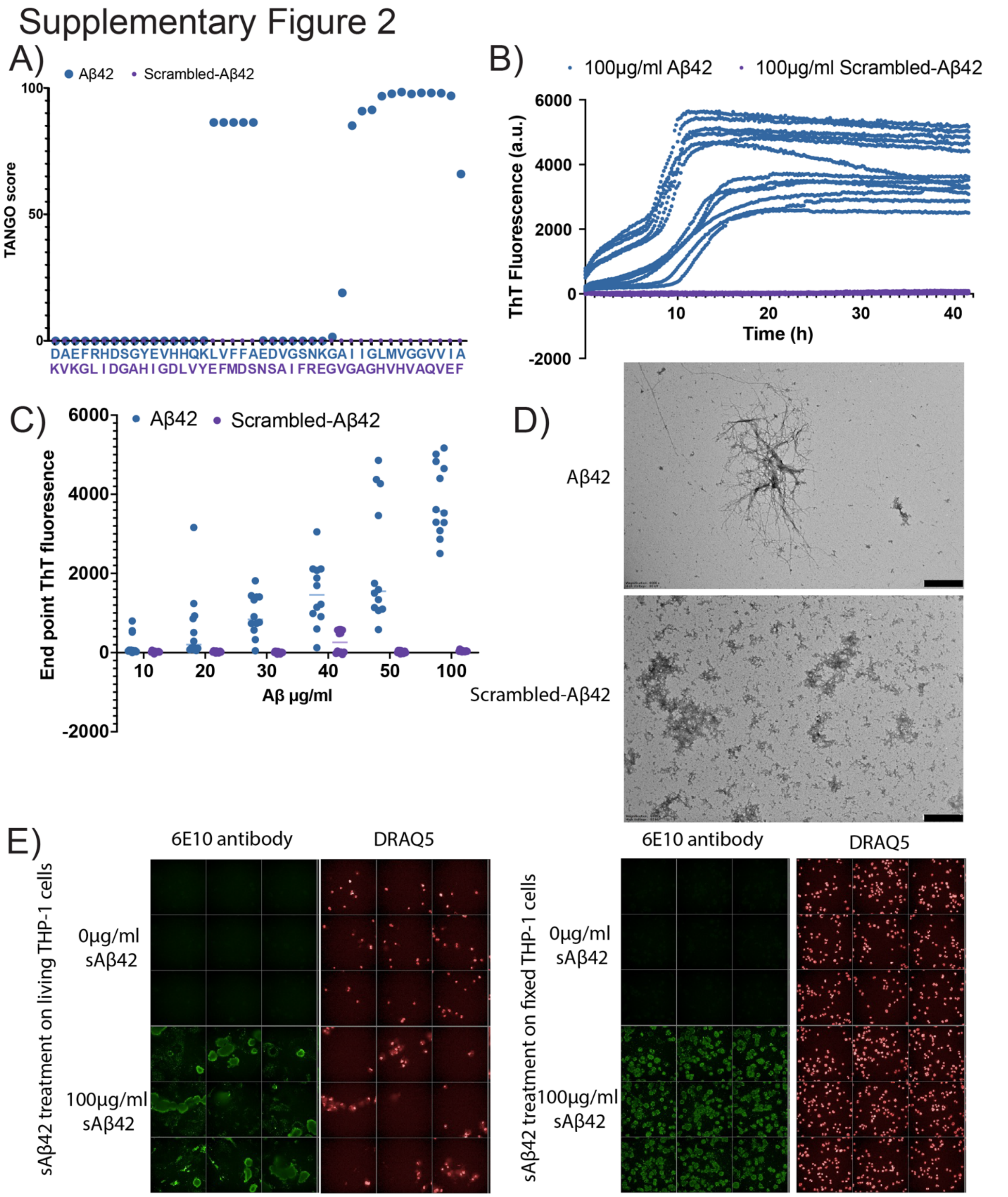


**Supplementary figure 2. Characterization of Scrambled-Aβ42**

A) TANGO score per amino acid of Scrambled-Aβ42(purple) and Aβ42(blue).

B) ThT kinetics of 100μg/ml Scrambled-Aβ42(purple) and Aβ42(blue). 2 independent experimets with 6 repeats.

C) End point fluoresence of different concentrations of Scrambled-Aβ42 (purple) and Aβ42 (blue).
2 independent experiments with 6 repeats.

D) TEM images of 100μg/ml Aβ42 and Scrambled-Aβ42. Scale bar: 1μm

E) Treatmet of living/fixed with sAβ42 and stained with 6E10 antibody(green) which recognize Aβ sequence and DRAQ5 (nuclei, red)
